## Supplemental_Figures for "Long non-coding RNAs direct the SWI/SNF complex to cell-specific enhancers"

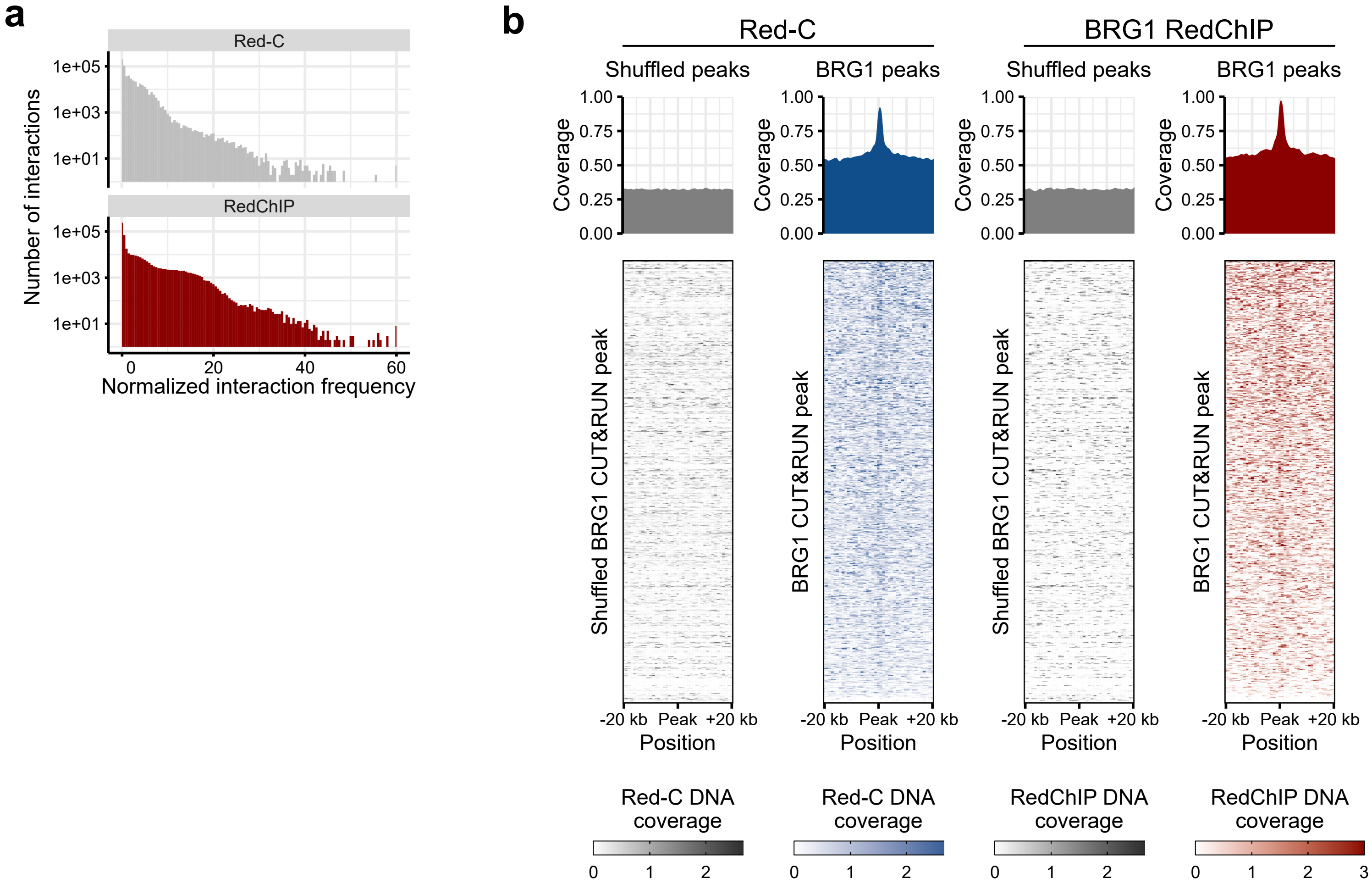


**Extended figure 1 | BRG1 RedChIP and CUT&RUN**

**(a)** Red-C and BRG1 RedChIP coverage. Number of RNA-DNA interactions across the normalized interaction frequencies

**(b)** Genome-wide density plots for RNA-DNA interactions for both Red-C and RedChIP DNA coverage at BRG1 CUT&RUN peaks versus a size-matched set of shuffled peaks

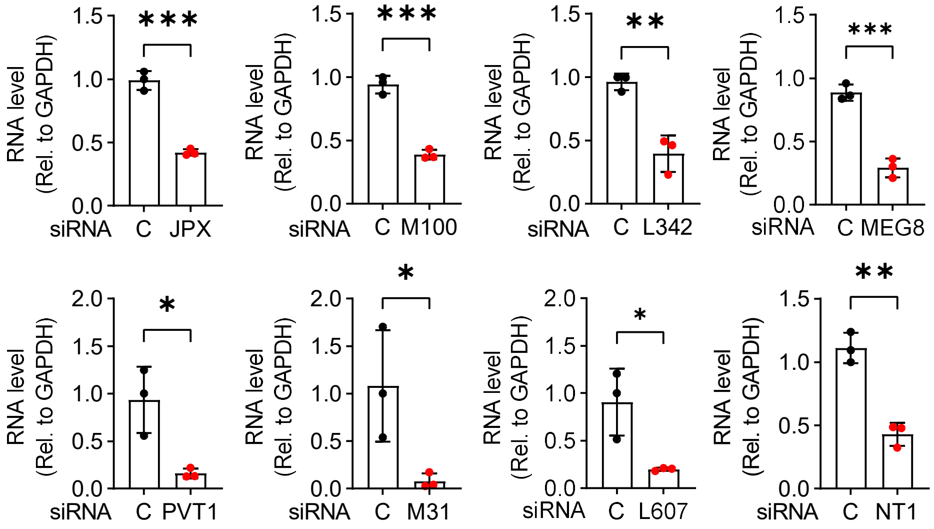


**Extended figure 2 | RT-qPCR validation of lncRNA candidate knockdowns**

siRNA against *JPX*, *MIR100HG* (M100), *LINC00342* (L342), *MEG8*, *PVT1*, *MIR31HG* (M31), *LINC00607* (L607) and *NEAT1* (NT1) followed by RT-qPCR. RNA level relative to GAPDH RNA. n=3, unpaired t-test. $p < 0.05$ indicated by *, $p < 0.01$ indicated by ** and $p < 0.001$ indicated by ***.

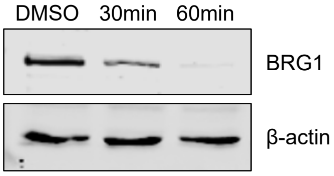


**Extended figure 3 | PROTAC AU-15330**

Western blot of HUVEC treated with 1 μM AU-15330 PROTAC (30 and 60 min) or DMSO (60 min). Antibodies against BRG1 and β-actin.
